# Mass Control Genes Buffer a Conserved Sublinear Scaling Law Linking Single-Cell Dry Mass to Mitochondrial Redox Efficiency

**DOI:** 10.64898/2026.04.18.719417

**Authors:** Zhaohang Li, Ziyue Yao, Shilin Ma, Jiali Su, Lewis C. Cantley, Ng Shyh-Chang

**Affiliations:** Key Laboratory of Organ Regeneration and Reconstruction, Institute of Zoology, Chinese Academy of Sciences, Beijing 100101, China; University of Chinese Academy of Sciences, Beijing 100049, China; Beijing Institute for Stem Cell and Regenerative Medicine, Beijing 100101, China; Dana-Farber Cancer Institute, Harvard Medical School, Boston, United States of America

**Keywords:** Single-cell mass, Cell growth, Cell metabolism, Mitochondrial redox, Scaling law

## Abstract

Using quantitative phase microscopy and genetically-encoded metabolism probes, we simultaneously measured single-cell dry mass and mitochondrial redox efficiency across eukaryotic cell types. We discovered that single-cell mass scales with mitochondrial redox efficiency, according to a sublinear power law, across budding yeast, mouse, and human cells. Genetic and pharmacological perturbations of mitochondrial redox balance shifted the coefficient *k* and exponent *y* predictably, while preserving the scaling law, indicating a causative coupling between biomass accrual and mitochondrial redox efficiency. We also discovered that both single-cell mass and mitochondrial redox efficiency manifest log-normal distributions, providing more support for previous theories that biomass accrual is an exponential process, meaning the rate of mass increase depends on the current mass. Finally, our yeast knockout screen identified 81 mass control genes (MCGs) that disrupted the log-normality of single-cell mass. Interestingly, MCGs were enriched for mitochondrial metabolism and cristae maintenance, whereas <10% were related to the cell cycle. Conserved MCGs, including human TRIAP1, ATPAF1, and ACO1, also maintained human single-cell mass’ log-normality and sublinear scaling with mitochondrial redox efficiency. These findings reveal a shared bioenergetic principle at the single-cell level, and new techniques to study it. This scaling law provides a quantitative framework for understanding cell mass control, metabolic coupling, and disease-associated growth regulation.

## Introduction

Quantitative scaling laws reveal some order within the random zoo of biological diversity. The observation that organismal metabolic rate scales sublinearly with body mass, has guided agriculture, ecology, pharmacology and physiology research for nearly two centuries. Proposed explanations for sublinear scaling invoke surface area-to-volume ratios, fractal transport networks, energy minimization, or criticality in nonequilibrium systems (West, Brown & Enquist, Science 1997; West et al., PNAS 2002; Yang et al., PNAS 2021), yet the single-cell origins of such allometric relationships remain unclear. If biomass accrual and bioenergetics are fundamentally linked at the organismal level, do analogous scaling principles govern the relationship between cell size and mitochondrial function?

Here, we combine quantitative phase microscopy for label-free single-cell dry mass measurement (scMass) with the mito-Grx1-roGFP2 reporter for mitochondrial glutathione redox potential, to test for any connection between cell mass and mitochondrial reactive oxygen species (mtROS) and redox efficiency. Across budding yeast, mouse, and human cell types, and spanning stem, primary, and cancer lineages, we find that scMass scales sublinearly with mitochondrial redox efficiency according to a conserved sublinear power law (exponent y ≈ 1/3). Precise genetic and pharmacological perturbations of mtROS shift the scaling parameters predictably while preserving the sublinear scaling with scMass, indicating that biomass accrual is causatively related to mitochondrial redox state. A yeast genetic loss-of-function screen identified 81 scMass-control genes (MCGs) that disrupted scMass log-normality and sublinear scaling, many of which are conserved in humans and enriched for mitochondrial metabolism and maintenance. These results reveal a fundamental bioenergetic principle that constrains the allowable metabolic state space at the single-cell level, with cell-type-specific exponents reflecting metabolic set-points. Given the fundamental interest in understanding cell size and cell growth, our findings have fundamental relevance for our understanding of biology (Ginzburg, Kafri & Kirschner, Science 2015).

## Results

Quantitative phase imaging refers to techniques that quantify the phase delay of a light wave of wavelength λ passing through a sample (Kim et al., 2014). The absolute dry mass of a single cell should be proportional to its DPC digital phase contrast value, multiplied by its area (Supplementary Equations), which can be automatically measured on standard microscopes or high-throughput imaging platforms (Figure 1A; Ali et al., 2008; Liu, Yan, Kirschner 2024).

**Figure 1.**
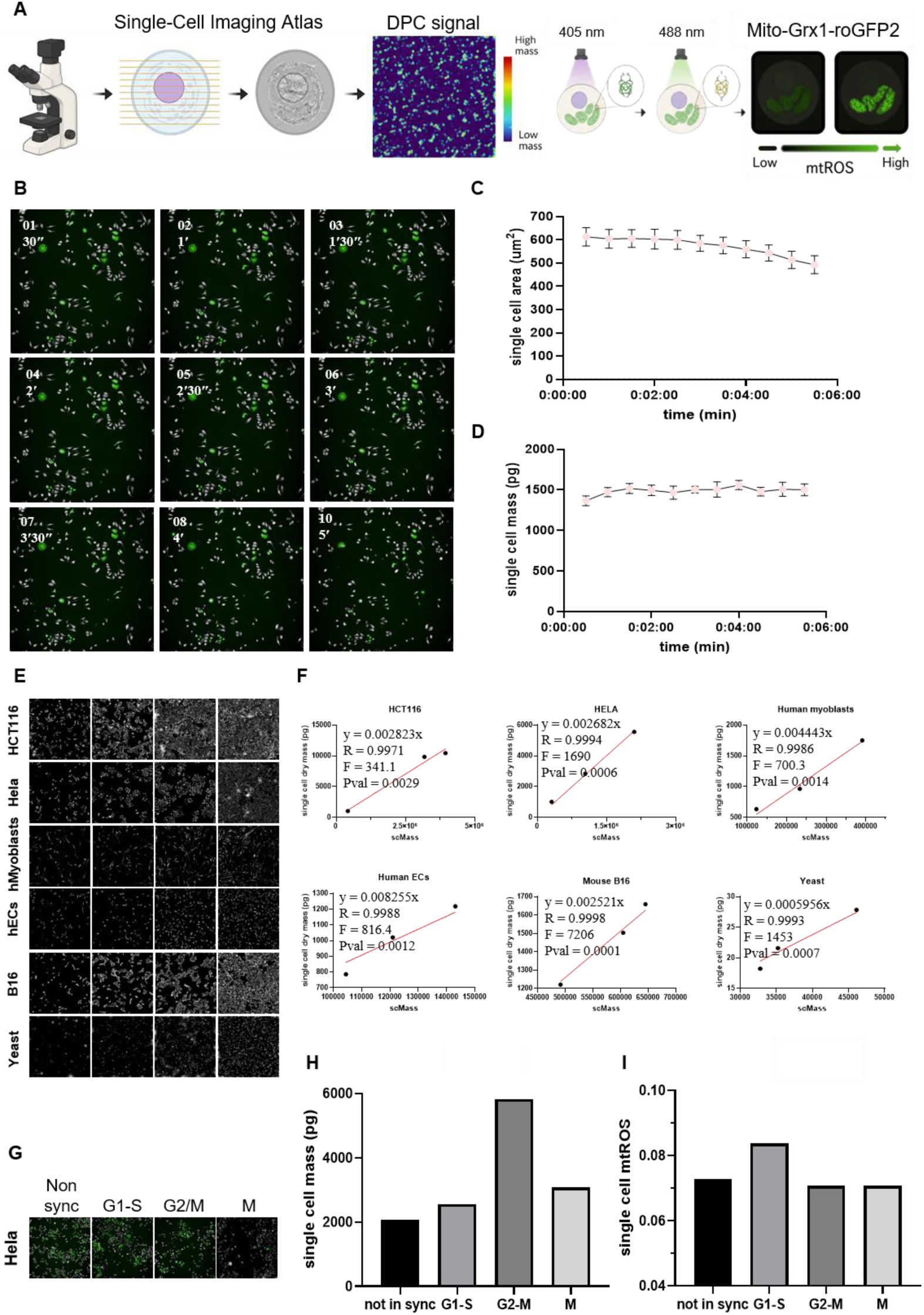
Using digital phase contrast (DPC) to measure single-cell mass (scMass), and the mito-Grx1-roGFP2 fusion protein reporter to measure mtROS. (A) Workflow schematic. (B) Representative DPC images and (C) Measurements of HeLa cells’ single cell areas and (D) single cell masses, measured by DPC, as they undergo trypsinization-induced shrinkage. (E) Representative DPC images and (F) Measurements of dry mass/cell of different cell-types at different growth densities is proportional to the sums of single cell mass (scMass) measured by DPC. (G) Representative images and (H) Live measurements of scMass, calibrated to dry mass, as the Hela cells undergo mitosis after nocodazole synchronization. (I) Live measurement of mtROS (i.e. mitochondrial glutathione redox potential) as HeLa cells undergo mitosis.

To test if our DPC-based derivation and measurements of scMass were robust and stable, we followed previously published methods (Liu et al., 2020) and subjected cells to trypsinization, and observed their single cell mass changes over time (Figure 1B). Theoretically, trypsinization would lead to rapid decreases in single cell areas (Figure 1C), but not affect single cell masses over a short time window before nutrient starvation effects manifest. Our results showed that simple DPC-based methods of single cell mass measurements were robust and stable to morphological changes (Figure 1D).

We further tested this derivation and DPC imaging of scMass by allowing a variety of cell-lines with different morphologies to grow to different densities, capturing live images for DPC and area analyses (Figure 1E), followed by harvesting for bulk dry mass measurements on a sensitive mass balance. Our results showed an excellent linearly proportional relationship between bulk dry mass and the area integral of single-cell DPC, across a variety of human (immortalized HCT116, HeLa, primary human skeletal myoblasts, primary endothelial cells), mouse (B16 melanoma) and yeast cells (Figure 1F). Using these linear calibration curves, we could image and calculate the absolute scMass of any cell-line using Eq 2b. One simple question we could address, to test the reliability of our measurements, would be how do single cells’ mass change at each stage of mitosis. To test this, we used nocodazole to synchronize Hela cells, which can reversibly induce cell cycle arrest at the G2/M checkpoint by reversibly inhibiting the formation of the mitotic spindle. In this way, we imaged and assessed scMass over the stages of mitosis from M to G1/S to G2/M, relative to their non-synchronized state, and calibrated by flow cytometry (Figure 1G). One could observe an increase in scMass as the cells progressed from the G1/S to the G2/M points (Figure 1H), as one might expect. The cells’ average scMass approximately halved at the M mitosis point from a peak at G2/M. Another interesting observation was that using this label-free method, we could assess that the non-synchronized Hela cells tended to be in the early G1 phase on average (Figure 1H), consistent with previous findings using fluorescent labels and flow cytometry (Ree et al., 2006).

Next, we used lentiviral infection of the mito-Grx1-roGFP2 fusion protein reporter at low MOI, to use its fluorescence ratio to measure mtROS levels (Gutscher et al., 2008; Supplementary Equations) in single cells. We assessed average mtROS levels at every stage of mitosis, and found that steady-state mtROS tended to be higher during the G1-S stage (Figure 1I), consistent with the higher metabolic rates needed to fuel biosynthesis during cell growth.

We then examined the overall DPC, area, mass and mtROS distributions of the single cells, and found that they all followed log-normal distributions (Figures 2A-D). Similar single cell mass average values and one log_10_ intervals in single cell mass variation (Figure 2C) have also been observed previously (Sims & Allbritton, 2007; Chen et al., 2020; Liu, Yan, Kirschner 2024). The appearance of log-normal distributions corroborates the idea that cell growth involves multiplicative biosynthetic processes between cell divisions, (Tzur et al., 2009; Turner, Ewald & Skotheim, 2012; Futcher & Kellogg, 2024), and a log-normal distribution naturally emerges from the central limit theorem applied to multiplicative variables with noise. To the best of our knowledge, we are the first to report log-normal distributions for single cell mtROS as well (Figure 2D).

**Figure 2.**
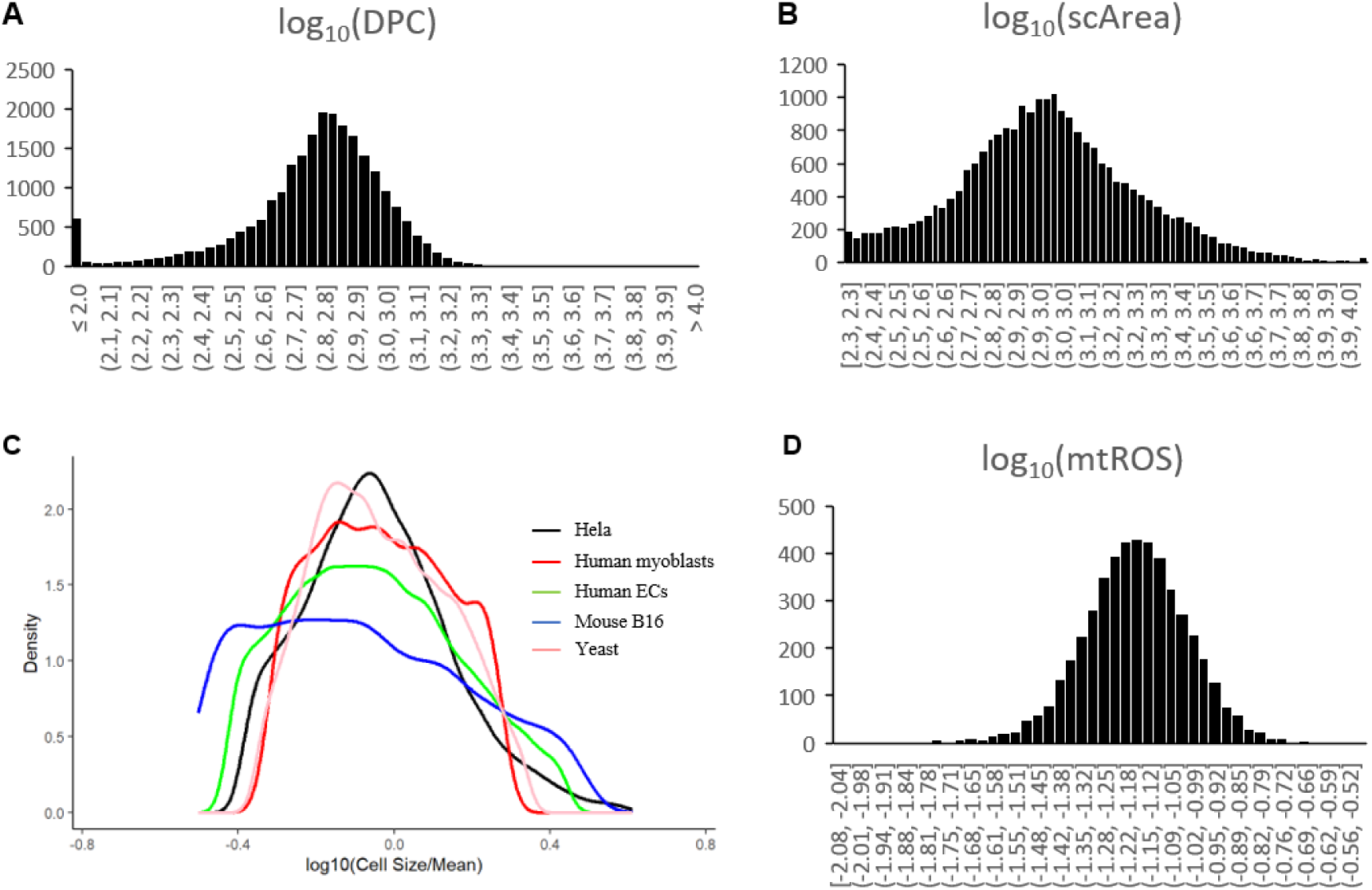
Single cells show log-normal distributions in mass and bioenergetic parameters. Log-normal frequency distributions of single Hela cells’ (A) DPC values, (B) areas, (C) renormalized dry mass (size/mean), relative to several cell-types, (D) mitochondrial reactive oxygen species (mtROS).

When examined in greater detail, to our surprise, we observed a negative relationship in log-log plots of scMass vs. mtROS, for every immortalized and primary cell-type we examined (Figures 3A-D). As expected, the immortalized cancer cells have higher average single cell masses, than the primary human cells and yeast cells, and it is known that across cell-types the mass and volume of cells can vary more than 1000-fold (Ginzburg, Kafri & Kirschner, 2015; Futcher & Kellogg, 2024). Despite the variation in scMass across cell-types, all relationships fit a sublinear power law (confidence P<0.001, power-law fit F-statistic>100, correlation R≈0.3), with exponents spanning *y* ≈ 0.2–0.4 (average≈0.3, Table 1). The exponent variation suggests that while the mass-mtROS scaling form is conserved, the precise mass-mtROS trade-off is tunable across cell-types. Altogether, the statistics confirm that mass-mtROS scaling is a highly consistent, population-level biological trend rather than a sampling artifact, and that mtROS is at least a partial contributor within a highly complex, multifactorial network for scMass regulation.

**Figure 3.**
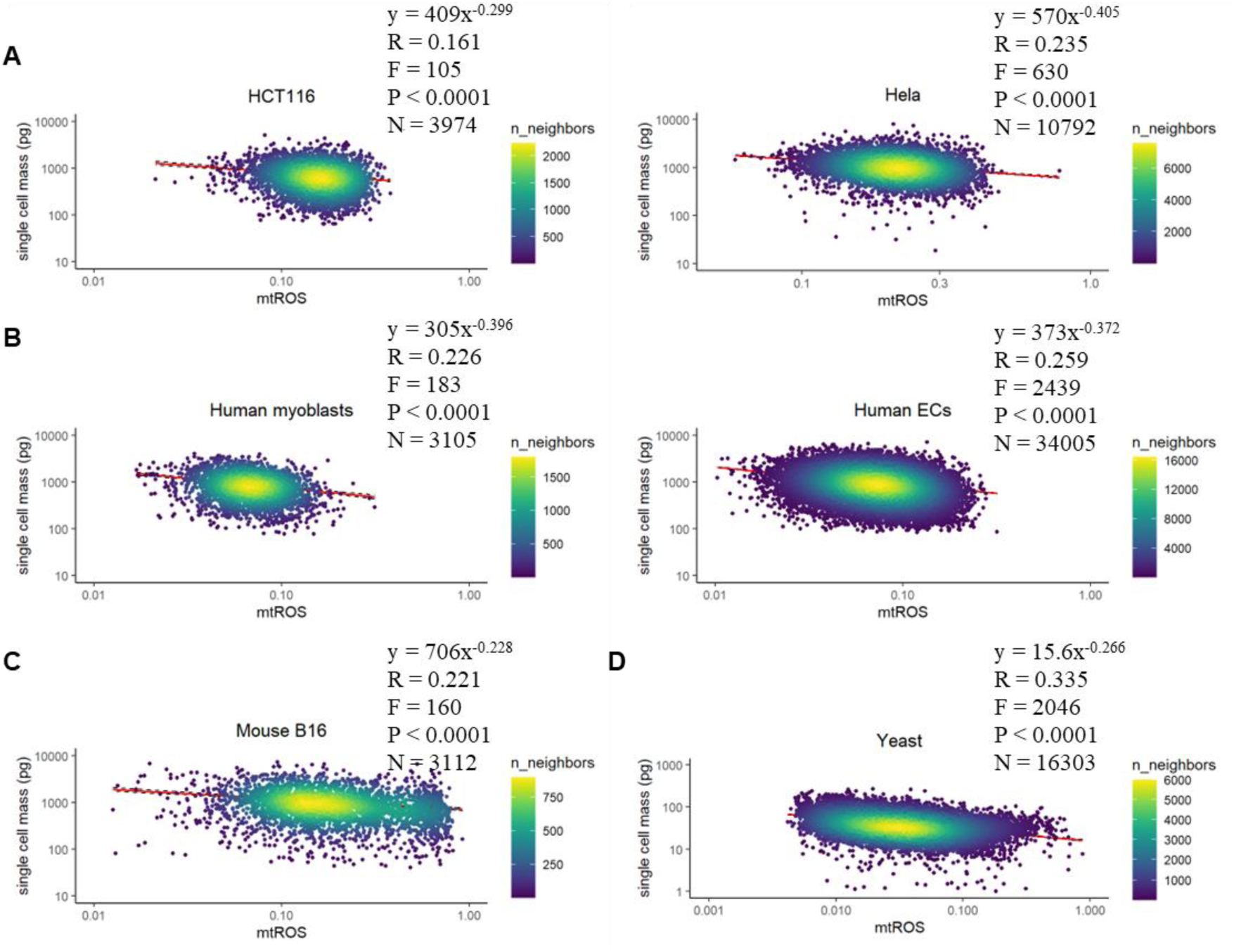
Single cell dry mass (pg) vs. mtROS fits a sublinear power law for many cell-lines, with a negative exponent ≈ 0.3. (A) Human cancer cells, HCT116 and Hela. (B) Human primary cells, skeletal myoblasts and endothelial cells (ECs). (C) Mouse B16 melanoma cells. (D) Budding yeast cells.

In comparison, when we examined the scMass vs mitochondrial NADH/NAD ratio, using the fluorescence ratio of another mitochondrial-localized GFP-based ratiometric reporter in single cells (Hu et al., 2021), no significant scaling relations were observed (P=0.431; Figure S1). These empirical observations suggest it might not be bioenergetics at large that is regulating biomass accrual, but there is a specific principle governing the scaling relationship between scMass and mitochondrial redox efficiency, regardless of cell-type and species.

However, correlation does not imply causality. To test the response of these exponents to antioxidants/ROS, and the functional dependence of scMass on mtROS, we subjected primary human myoblasts to treatment with the general antioxidant N-acetyl-cysteine (NAC) as well as the mtROS-inducing drug rotenone, a mitochondrial Complex I-specific inhibitor that does increase average mtROS as expected (Figures S2, S3A).

We found that compared to the DMSO vehicle control, NAC antioxidant treatment led to a lower *y* (from 0.336 decreased to 0.214) and a higher *k* (from 302 increased to 394; Figures 4A-C). If scMass was dependent on OxPhos-generated mtROS, we would expect that the mtROS-inducing drug rotenone would have the opposite effect of NAC antioxidant. However, if scMass was merely dependent on OxPhos-generated ATP, we would expect that rotenone (lower ATP too) would have the same effect as NAC (lower ATP too, due to energy-depleting glutathione biosynthesis from NAC, Figure S2). Of course, scMass could also be independent of rotenone or Complex I, and thus not change after treatment. We found that compared to the DMSO vehicle control, rotenone treatment led to a higher *y* (from 0.336 increased to 0.362) and a lower *k* (decreased to 297; Figures 4A-C), indicating that scMass is indeed dependent on mtROS.

**Figure 4.**
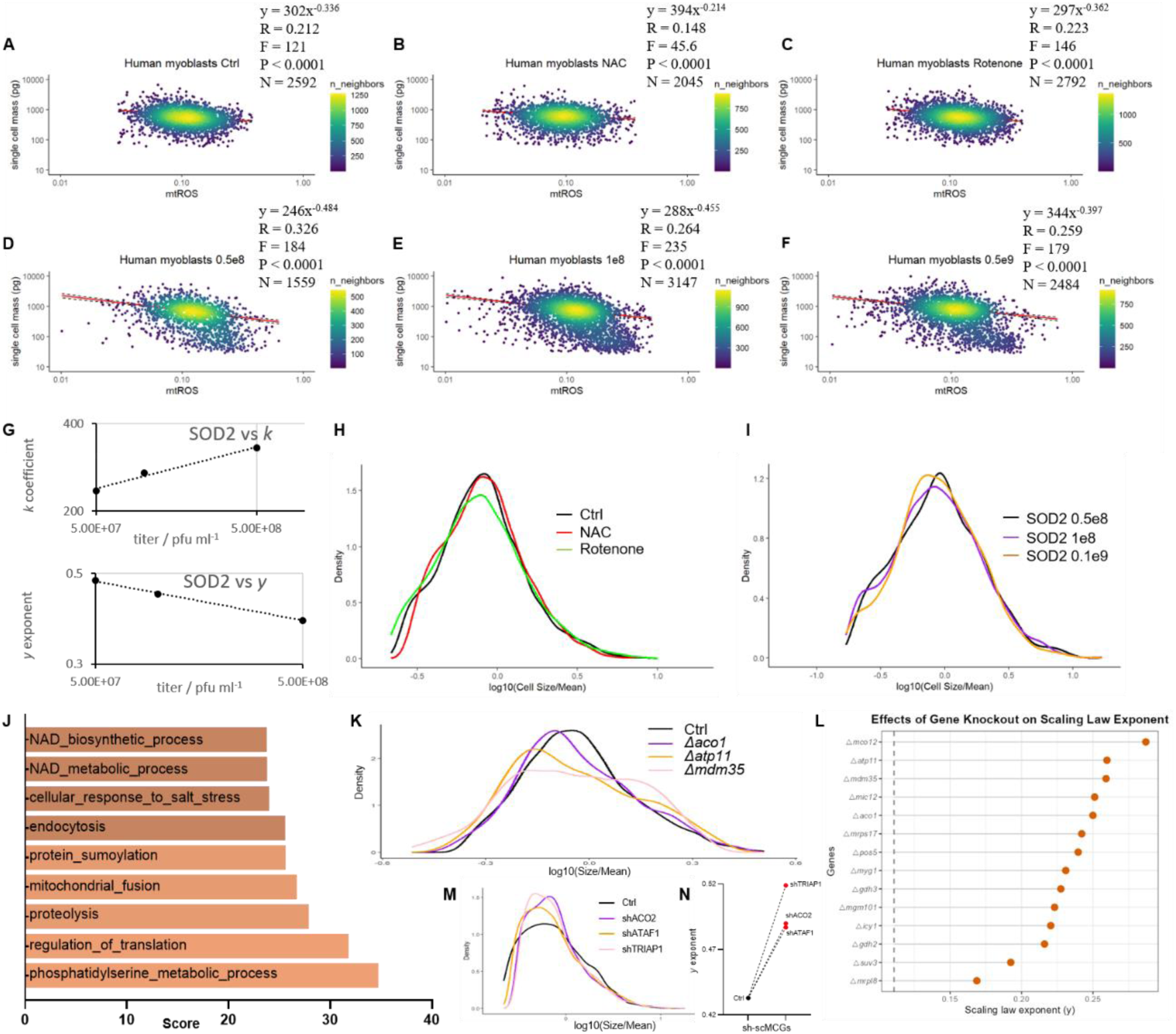
Effects of mtROS perturbation on scMass-mtROS scaling law. (A) Single cell mass (pg) vs mtROS in primary human skeletal myoblasts treated with the vehicle control DMSO, (B) 20mM N-acetyl-cysteine (NAC), or (C) 50nM Rotenone, for 24 hours. (D) Single cell mass (pg) vs mtROS in primary human skeletal myoblasts infected with adenovirus containing SOD2-2A-mitoCAT, at the dosages of 0.5×10^8^, (E) 1.0×10^8^, or (F) 0.5×10^9^ pfu/ml. (G) Plots of SOD2-2A-mitoCAT adenovirus dosages vs coefficients k and exponents y. (H) Log-normal distribution of single-cell masses in human myoblasts treated with DMSO, NAC and Rotenone. (I) Log-normal distribution of single-cell masses in human myoblasts infected with different doses of SOD2-2A-mitoCAT adenovirus. (J) Gene Ontology enrichment analysis of MCGs identified from the Dharmacon YKO budding yeast knockout library. (K) Effects of MCG knockout on yeast cells’ scMass log-normal distribution. (L) Effects of MCG knockout on the increase in yeast cells’ scaling exponent y. (M) Effects of MCG knockdown on myoblasts’ scMass log-normal distribution (N) Effects of MCG knockdown on myoblasts’ scaling law exponent y.

To further confirm these findings, we used an orthogonal set of mtROS-specific, non-bioenergetic antioxidant enzymes: mitochondria-specific superoxide dismutase 2 (SOD2) and mitochondria-targeted catalase (mitoCAT). We performed titrations of SOD2-2A-mitoCAT adenovirus to steadily and precisely decrease mtROS (Figure S3B). If the conclusions from the pharmacological experiments were true, we would expect increasing titers of SOD2-2A-mitoCAT to steadily decrease mtROS like NAC and thus decrease *y*, while increasing *k*. Indeed, we found that increasing the SOD2-2A-mitoCAT adenovirus titer from 5×10^7^ to 1×10^8^ to 5×10^8^ pfu ml^-1^ (Figures 4D-F), led to a steady increase of *k* from 246 to 288 to 344 while *y* steadily decreased from 0.484 to 0.455 to 0.397 (Figure 4G). We also noticed that, while DMSO did not significantly affect the baseline coefficient *k* and exponent *y* of primary human myoblasts, adenovirus titer does affect the baseline coefficient *k* and exponent *y* of primary human myoblasts (see Table 1), suggesting that viral protein production or antiviral defenses can affect the OxPhos production of mtROS. However, none of these perturbations fundamentally affected the overall sublinearity of the scaling law scMass = *k*[mtROS]^-*y*^, nor the overall log-normality of single cell masses (Figures 4H, I), indicating the robustness of scMass-mtROS scaling.

**Table 1.** Multiple cell-types have a characteristic scMass-mtROS scaling law.

| **Cell-type** | **Exponent $y$** | **Coefficient $k$** |
| --- | --- | --- |
| HEK293T | 0.260 ± 0.003 | 682 ± 3.1 |
| HCT116 | 0.299 ± 0.005 | 409 ± 3.8 |
| Hela | 0.405 ± 0.002 | 570 ± 1.4 |
| Primary Human myoblasts | 0.396 ± 0.003 | 305 ± 2.3 |
| Primary Human EC | 0.372 ± 0.007 | 373 ± 5.0 |
| Mouse B16 | 0.228 ± 0.004 | 706 ± 1.7 |
| Yeast | 0.266 ± 0.09 | 15.6 ± 0.7 |

Thus, both pharmacological and genetically precise mtROS perturbations demonstrate that the scaling parameters (*k, y*) are causally dependent on mtROS. Lowering mtROS consistently decreases *y* and increases *k*, while increasing mtROS shifts the scaling parameters in the opposite direction. Critically, none of these perturbations abolish the sublinear scaling relation, indicating that the coupling is a consistent feature of eukaryotic metabolism. The exponent range (y ≈ 0.2–0.4) thus represents a bounded, tolerable parameter space for metabolic set-points, rather than a fixed physical constant.

To delve into the molecular origins of this scaling relation, we leveraged the power of yeast genetics to rapidly screen and identify genes that affect the log-normality of scMass distributions, which would necessarily affect the exponents of the scaling relations. We named these Mass Control Genes (MCGs). We identified only 81 MCGs, out of a total of over 6000 genes in the complete yeast knockout library, that were required to maintain the log-normality of scMass (Table S1). These MCGs were significantly enriched for mitochondrial and metabolism genes, according to Gene Ontology enrichment analyses (Figure 4J). Thus, we selected 15 mitochondria-related MCG knockout strains to transform with the mito-Grx1-roGFP2 reporter to test the effects of MCG knockout on the scaling law. Interestingly, as these yeast knockout strains’ scMass distributions showed deviations from log-normality, all of their sublinear exponents also increased (Figures 4K,L), implying the knockout of mitochondria-related MCGs led to the loss of cell mass regulation and then a general shift from sublinear towards linear scaling. It appears that we have identified a new network of mitochondria-related MCGs for maintaining scMass in a log-normal distribution, using negative feedback loops to buffer the effects of noisy mtROS on cell growth self-proportionality within a sublinear scaling law.

To verify if some of these mitochondria-related MCGs are conserved in human cells, we selected the top hits and performed shRNA knockdowns in human myoblasts. We found that the human mitochondrial cristae phosphatidylethanolamine transporter *TRIAP1* (yeast *mdm35*), the human mitochondrial ATP synthase chaperone *ATPAF1* (yeast *atp11*), and the human mitochondrial aconitase *ACO2* (yeast *aco1*) were also conserved MCGs. Their knockdown led the scMass distributions to show deviations from log-normality (Figure 4M) and all the sublinear exponents to increase towards linearity (Figure 4N), consistent with their yeast counterparts. Mitochondrial aconitase (yeast *aco1*) uses its [4Fe-4S] cluster, one of the most mtROS-sensitive enzymatic sites in the mitochondrial matrix, to buffer against the effects of mtROS on citrate flux into growth-related metabolic pathways. TRIAP1 (yeast *mdm35*) controls mitochondrial phosphatidylethanolamine transport to the inner membrane to maintain cristae curvature and ETC supercomplex assembly, thus buffering against the effects of mtROS and phospholipid peroxidation on mitochondrial bioenergetics and cell growth. ATPAF1 (yeast *atp11*) is a chaperone for F1-ATP synthase assembly by binding to F1 α and β subunits post-translationally, thus buffering against mtROS by dissipating the mitochondrial membrane potential with ATP synthase. As a sign of their specificity, these genes’ paralogs and most of the other mitochondrial genes in the yeast whole-genome knockout library were not in the list of MCGs, providing yet more internal controls for the yeast genetic screen.

Given the fact that mito-Grx1-roGFP2 and DPC signals were obtained in asynchronous yeast populations, we further evaluated the influence of the cell cycle on the scaling law. Among the 81 MCGs identified, only 4 genes have known cell-cycle-related phenotypes: *Δkin4* causes faster mitotic exit, whereas *Δcsm1* causes a metaphase delay, and *Δrad27/rpa12* causes a S-phase delay. However, the fact that only ∼5% of the 81 MCGs have known cell cycle phenotypes, meant that cell cycle disruption is not the only driving factor for the changes we observed in the yeast knockout screen. Moreover, we observed that even amongst synchronized cells at the same cell cycle phase, the same sublinear scaling law was still observed (Figure S4).

## Discussion

Allometric scaling laws have shaped how we think about biology across levels of organization for two centuries. Here we show that a sublinear power law connects single-cell dry mass to steady-state mitochondrial ROS across eukaryotic cell types. This negative power law (scMass = *k*[mtROS]⁻*ʸ*) holds from yeast to human cells, with exponents spanning *y* ≈ 0.2–0.4. This range likely reflects cell-type-specific metabolic set points, whereby cancer lines, primary cells, and yeast occupy distinct positions along a shared biophysical principle.

Recent work suggests that certain specific metabolic genes and pathways can directly constrain and regulate cell growth (Shyh-Chang, Daley, Cantley 2013; Talavera et al., 2024). Our yeast screen further identified 81 MCGs that tighten the log-normality of scMass and stabilize the scaling exponent’s sublinearity (prevent excessive scaling). These MCGs are enriched for mitochondrial functions, including cristae maintenance, ATP synthase assembly, redox-sensitive metabolism. Mitochondrial aconitase provides a concrete example. Its mtROS-sensitive [4Fe-4S] cluster senses and couples mtROS to citrate-lipid anabolic flux vs. isocitrate-TCA-mtROS catabolic flux with negative feedback loops (Tong & Rouault, 2007), although the magnitude of this feedback loop may be variable depending on the cell-type’s metabolic set points. The mitochondrial MCG network appears to constrain this variability, preventing extreme exponent values that would destabilize growth. Notably, we recovered no mitochondrial MCG knockouts that flatten or abolish the scaling law, we only found knockouts that accentuate the scaling slope. This suggests that these mitochondrial MCGs are actively buffering the scaling exponent within a sublinear range. Seen from this perspective, these newly identified mitochondrial MCGs would constitute a new mitochondrial gene network to buffer a fundamental scaling law between cellular biomass accrual and mitochondrial redox efficiency.

Pharmacological and genetic perturbations that alter mtROS shift both scaling parameters (*k, y*) predictably while preserving a negative exponent, indicating a causal relationship with parameter tunability. The negative sublinear power law persists through these perturbations, suggesting the coupling is a consistent feature of eukaryotic biology. The exponent range likely emerges from heterogeneity in mitochondrial redox efficiency across cell types. In this study, mitochondrial redox <u>in</u>efficiency can be defined as the moles of mtROS produced per mole of oxygen consumed. Thus, higher mitochondrial redox efficiency can indicate a greater proportion of oxygen consumption rate (OCR) is directed towards ATP production, rather than mtROS generation via electron leakage at ETC Complexes I/III or the grid network of mitochondrial FAD-dependent flavoproteins (e.g. fatty acid oxidation-coupled ETFDH, tricarboxylic acid-coupled αKGDH/PDH/mG3PDH, amino acid-coupled PRODH/DHODH, CHDH, all of which are also minor, non-ETC sources of mtROS). Hence, our scaling law is coupled to mtROS and not any particular source of mtROS. This multi-source perspective, and the regulatory complexity of NADH/NAD enzyme binding vs. probe measurement precision, could explain why the mitochondrial NADH/NAD ratio did not manifest scaling relationships with scMass, important as it is to mitochondrial redox efficiency. Indeed, previous studies had shown that fluorescence lifetime imaging (FLIM) of free NADH can produce images of relative NADH abundance, thereby inferring relative ETC flux and OCR at subcellular resolution (Yang et al., 2021), with implications for our understanding of cellular metabolic scaling. Nevertheless, regardless of the heterogeneous set points in mitochondrial redox fluxes, our study indicates that scMass scales <u>negatively</u> with mitochondrial redox <u>in</u>efficiency, and thus scales <u>positively</u> with mitochondrial redox efficiency (Supplementary Equations), subject to sublinear buffering by MCGs.

While both our scMass-redox efficiency scaling law and the Kleiber Law are sublinear scaling laws, the scMass-mtROS scaling exponent (*y* ≈ 0.2 – 0.4) differs from the Kleiber Law’s bioenergetics-biomass exponent (∼3/4). In fact, if we substitute for mass, we obtain the invariant equation K = [mtROS] ∗ OCR^4/3*y*^ (Supplementary Equations), i.e. low OCR correlates with high mtROS (and vice versa), which is well-known to hold true, because hypoxia causes electron flow to slow down in the ETC, highly reducing the CoQ pool, such that the probability of an electron leaking to form mtROS rises. Thus, the two sublinear scaling laws may reflect mutually-interacting constraints imposed by energy transduction kinetics and energy transmission geometry. At the cellular level, their exponents could reflect how mitochondrial membrane architecture, ETC supercomplexes, and redox regulation interact to scale with scMass. Testing this will require combining our imaging approach with ultrastructural and metabolic measurements. Nevertheless, while the current study pertains to the single-cell level, metabolic scaling at the organismal level is the result of contributions from many heterogeneous cell types and many tissue systems, such as the muscular, nervous and endocrine systems, where tissue-level heterogeneity in growth/metabolic rates cannot be ignored (Glazier, 2015). Therefore, at this stage, the two sublinear scaling laws may be similar only at the semantic level of relating mass to bioenergetics-related parameters. Future work is needed to relate metabolic scaling at the single cell level to metabolic scaling at the organismal level.

Although previous studies (Lanz et al., Mol Cell 2022) observed that mitochondrial proteins super-scale with cell volume, which ostensibly contradicts our sublinear scaling law, this super-scaling could be simply explained as a rearrangement in our equation. Sublinear scaling of scMass with mtROS, implies mitochondrial redox efficiency super-scaling with scMass (Supplementary Equations). Taken together, larger cells may have more mitochondrial mass, but their more advanced mitochondrial networks also leak fewer electrons to produce less mtROS per unit respiratory flux.

In a related study, Miettinen and Björklund (Dev Cell 2016; Fig 1b) displayed a negative linear relationship between mitochondrial membrane potential vs. cell size/volume in Drosophila cells, although they did not study it in detail. Instead, they focused on mass-normalized mitochondrial membrane potential and how it presents a bell-shaped curve with scVolume. While previous work highlighted cell-level optimality through a bell-shaped distribution of mass-normalized mitochondrial potential, our findings extend this observation by demonstrating the log-normality of both scMass and mtROS, and reveals that the underlying trade-off between biomass accrual and redox efficiency is governed by a robust, conserved sublinear power law in many eukaryotic cell types. In future work, it would be interesting to delve into the relationships between scMass and scVolume, and their relationships to cell cycle duration, polyploidy, development and senescence (Cadart et al., 2018; Cadart & Heald, 2022; Cadart et al., 2023; Miyazawa & Aulehla, 2018; Neurohr et al., 2019).

We conclude that our finding on the coupling between single-cell dry mass and mitochondrial redox efficiency provides a new framework to study all eukaryotes. This quantitative principle reframes cell mass control as an important and emergent property deserving of deeper studies.

## Supplementary Equations and Figures

### DPC to scMass

DPC (digital phase contrast) is proportional to the phase delay Δφ introduced by a cell, which is typically measured relative to the phase delay in the cell-free area. Thus, Δφ is proportional both to cell thickness *h* and to the refractive index (*n*) difference between the cell and surrounding liquid:

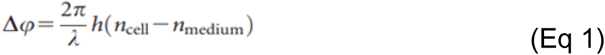

The column-averaged refractive index difference (Δ*n*_average_ *= n*_cell_-*n*_medium_) is equal to the intracellular density multiplied by the specific refractive increment constant (Theisen et al., 2000), and the product of intracellular density and cell volume yields the total dry mass. Thus, the surface area integral of each single cell’s phase delay Δφ (over area A) reflects the single cell’s total dry mass T (pg):

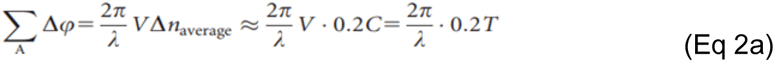

where λ is the illumination wavelength (μm), V is the cell volume (μm^3^), C is the specific refractive increment (for almost all eukaryotic cells, the constant C is remarkably stable at approximately 0.2 μm^3^/pg for biomacromolecules, e.g. proteins and nucleic acids). For a single cell under the microscope, multiplying its average Δφ with its area A and a geometry-specific coefficient *r* should allow us to obtain a discrete approximation of this area integral and thus the single cell’s dry mass T:

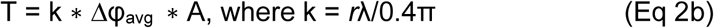

Since Δφ_avg_ is proportional to the DPC signal, the total dry mass of a single cell should be proportional to DPC ∗ A, and thus absolute scMass can be calibrated and measured on microscopes or high-throughput imaging platforms capable of single-cell DPC and area quantification (Ali et al., 2008).

### Mito-Grx1-roGFP2 to mitochondrial redox efficiency

Fusion of a mitochondria-targeted Glutaredoxin 1 (Grx1) to roGFP2 couples the roGFP2 probe’s redox state (roGFP2_red_ vs roGFP2_ox_) to the mitochondrial glutathione pool via a deglutathionylation/glutathionylation cycle. The reversible reaction is:

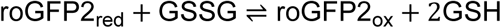

By the law of mass action, the equilibrium constant for this reaction is:

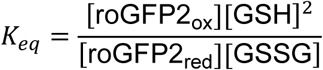

Rearranging this to solve for the ratio of oxidized to reduced probe yields:

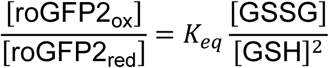

Our observable variable is the roGFP2 probe’s dual fluorescence signal ratio 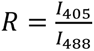.

The fraction of oxidized probe (*f_ox_*) is related to *R* by the dynamic range of the probe (*R_min_* for fully reduced, *R_max_* for fully oxidized):

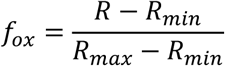

The ratio of oxidized to reduced probe is therefore a renormalized *R* :

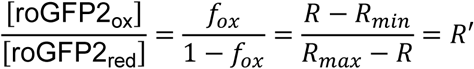

Equating the thermodynamic and optical expressions gives:

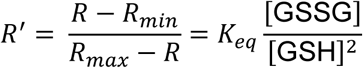

The equation above describes a thermodynamic equilibrium, but the cell is in a non-equilibrium steady state driven by mtROS production and glutathione enzymes. Although the mitochondrial glutathione pool is maintained at a non-equilibrium steady state by the opposing fluxes of glutathione peroxidase (GSSG production) and glutathione reductase (GSSG reduction), the mito-Grx1-roGFP2 probe reports the instantaneous steady-state ratio [GSSG]/[GSH]^2^ in mitochondria under the following conditions: (1) Grx1 catalyzes probe equilibration on a timescale (milliseconds) much faster than glutathione pool turnover (seconds to minutes), ensuring the probe remains in quasi-equilibrium with the glutathione pool; (2) Because [GSH] ≫ [GSSG] under physiological conditions (typically 100:1 to 1000:1), and the probe concentration is low relative to the glutathione pool, this ensures the probe’s redox effects do not perturb the mitochondrial redox system. Under these two conditions, the equilibrium-derived equation remains a valid reporter of the steady-state glutathione redox potential.

Mitochondrial ROS (e.g., *H*_2_*O*_2_) is scavenged by glutathione peroxidase (GPx). Because physiological [*GSH*](1–10 mM) vastly exceeds the *K_m_* of GPx for GSH, the rate of GSSG production is essentially independent of [*GSH*] and proportional only to mtROS:

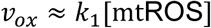

GR reduces GSSG back to GSH. Because physiological [*GSSG*] is typically ≪ *K_m_* of GR, the reduction rate is first-order:

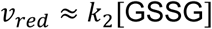

At steady state, we set *v_ox_* = *v_red_* to yield [GSSG]*_ss_* ∝ [mtROS].

Because the *square* of [*GSH*] is so large relative to [mtROS], it behaves like a constant in the same cell-type. Thus, substituting this back into the probe equation, we obtain:

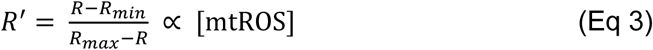

Thus, the mito-Grx1-roGFP2 probe’s normalized fluorescence ratio is approximately proportional to [mtROS] in the same cell-type.

In addition, if we define mitochondrial redox <u>in</u>efficiency as the number of mtROS (superoxide, peroxide, or hydroxyl radicals etc) moles generated per mole of ATP generated, then mitochondrial redox efficiency would be the inverse of this ratio. Assuming constant ATP synthesis rate at steady state, we can renormalize and approximate mitochondrial redox efficiency using *k*/*R’*, or −log(*R’*) on a log-log plot. Thus, we would obtain a left-right mirror image of the current scMass-mtROS scaling law, i.e. a positive sublinear scaling law for scMass vs. mitochondrial redox efficiency.

### scMass-mtROS scaling law vs. Kleiber’s Law

Kleiber’s Law (West et al., 2002): Single cell basal metabolic rate or *B* = k_1_M^3/4^
Our scMass law suggests: *B* = k_1_M^3/4^ = k_2_[mtROS]*^-y^*)^3/4^ (∼ k_2_[mtROS]^-1/4^ if *y*∼1/3)
Rearranged: K = [mtROS] ∗ B^4/3y^ (∼ [mtROS] ∗ B^4^ if *y*∼1/3)

As a result of Thornton’s Rule (Ilker et al., 2026), we can interconvert basal metabolic rate *B* and oxygen consumption rate OCR via a constant Δh_ox_ ≈ −450 kJ/mol, meaning:

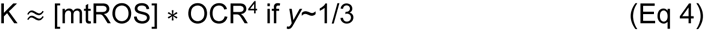

The equation suggests that mitochondrial ETC, and its associated grid of mitochondrial flavoproteins, can be modeled as a kinetically bifurcated pathway, reflecting a trade-off between bioenergetic flux and mtROS production flux. During transitions to hypoxia, [mtROS] would increase, and vice versa.

**Figure S1.**
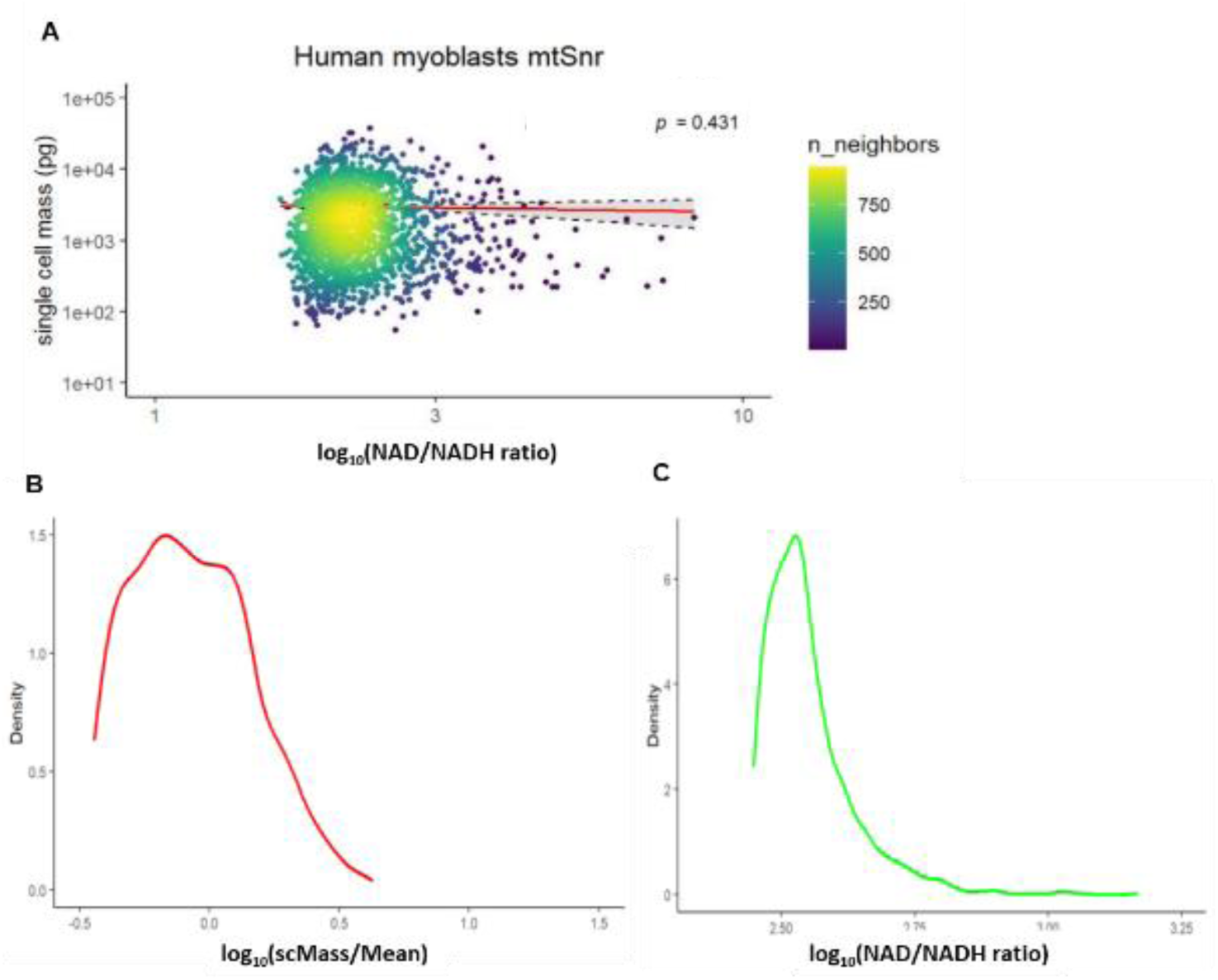
Single cell dry mass (pg) vs mitochondrial NAD/NADH ratio, does not fit a scaling law. (A) Single cell mass (pg) vs mitochondrial NAD/NADH ratio, as determined by a mitochondrial-localized SoNar reporter (mtSnr) in primary human skeletal myoblasts. (B) Log-normal distribution of single-cell masses in human myoblasts with mtSnr. (C) Log-normal distribution of single cell mitochondrial NAD/NADH ratios in human myoblasts with mito-SoNar.

**Figure S2.**
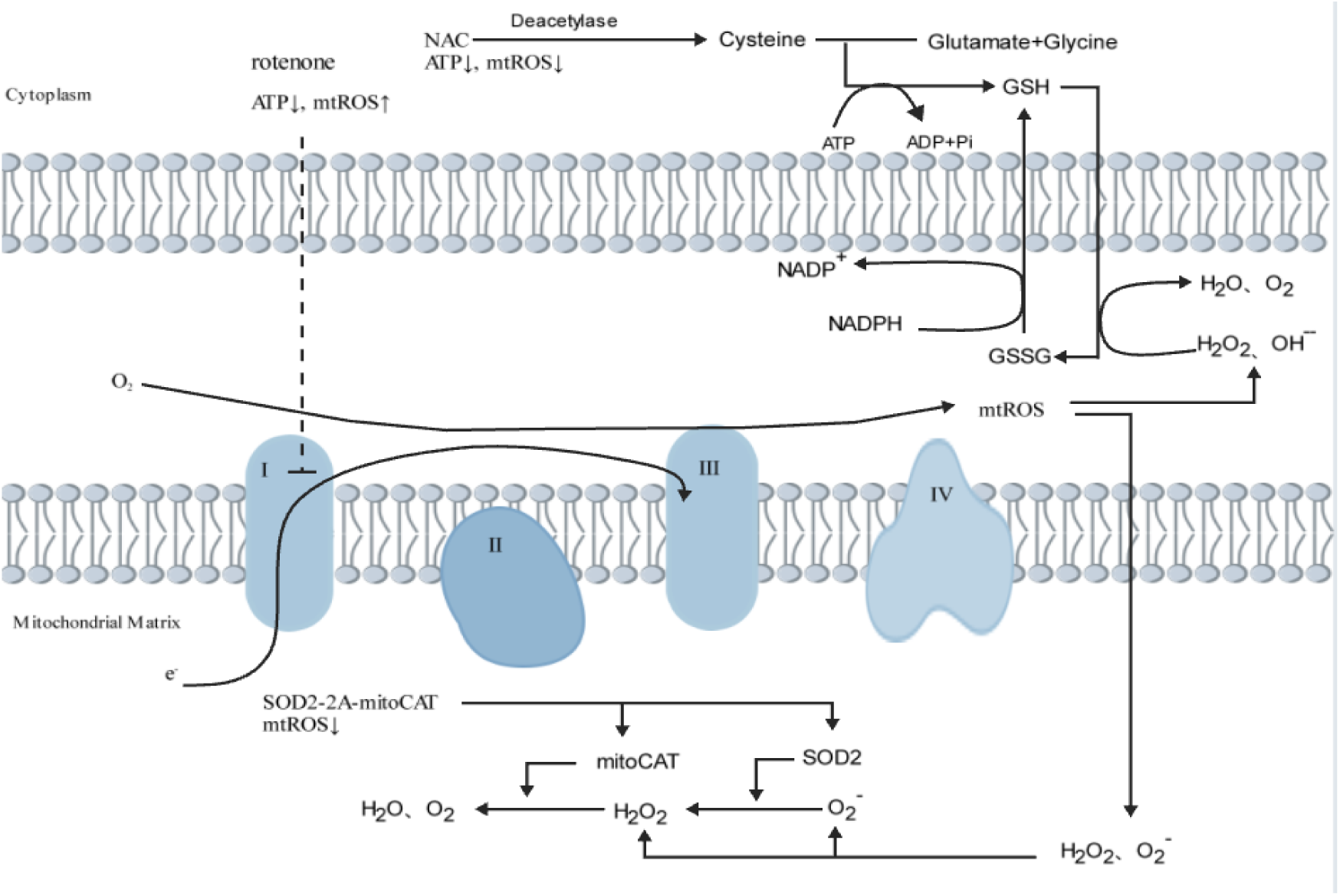
Schematic of mitochondrial ROS generation and modulation at the electron transport chain. Complex I–IV are embedded in the inner mitochondrial membrane. When the high energy electrons of NADH are gradually passed through Complexes I-IV of the electron transport chain (before being accepted finally by the highly electronegative O_2_ molecules to form H_2_O), the energy released by the "downhill" movement of electrons is coupled to the Complexes’ intermembrane transport of protons against a concentration gradient. This creates the electrochemical gradient, often called the proton-motive force, that powers ATP synthase. However, not every electron is passed through the Complexes smoothly, and electron leakage can lead to generation of mitochondrial reactive oxygen species (mtROS) such as superoxide anions O_2_-. Thus, for every O_2_-generated occasionally by inefficient electron transport, many more H_2_O are generated. Rotenone inhibits the electron transfer efficiency of Complex I, causing reverse electron flow, mtROS↑ and ATP↓. After a series of ATP-consuming enzymatic reactions, N-acetyl-cysteine (NAC) rapidly diffuses across plasma membranes and fuels the generation of glutathione (GSH) which is an antioxidant, resulting in ATP↓ and mtROS↓. SOD2-2A- mitoCAT is a transgene that generates two mitochondrial-localized antioxidant enzymes, whereby superoxide dismutase 2 (SOD2) converts superoxide O_2_- into peroxide H_2_O_2_ and mitochondrial-targeted catalase (mitoCAT) converts H_2_O_2_ to H_2_O, leading to mtROS↓.

**Figure S3.**
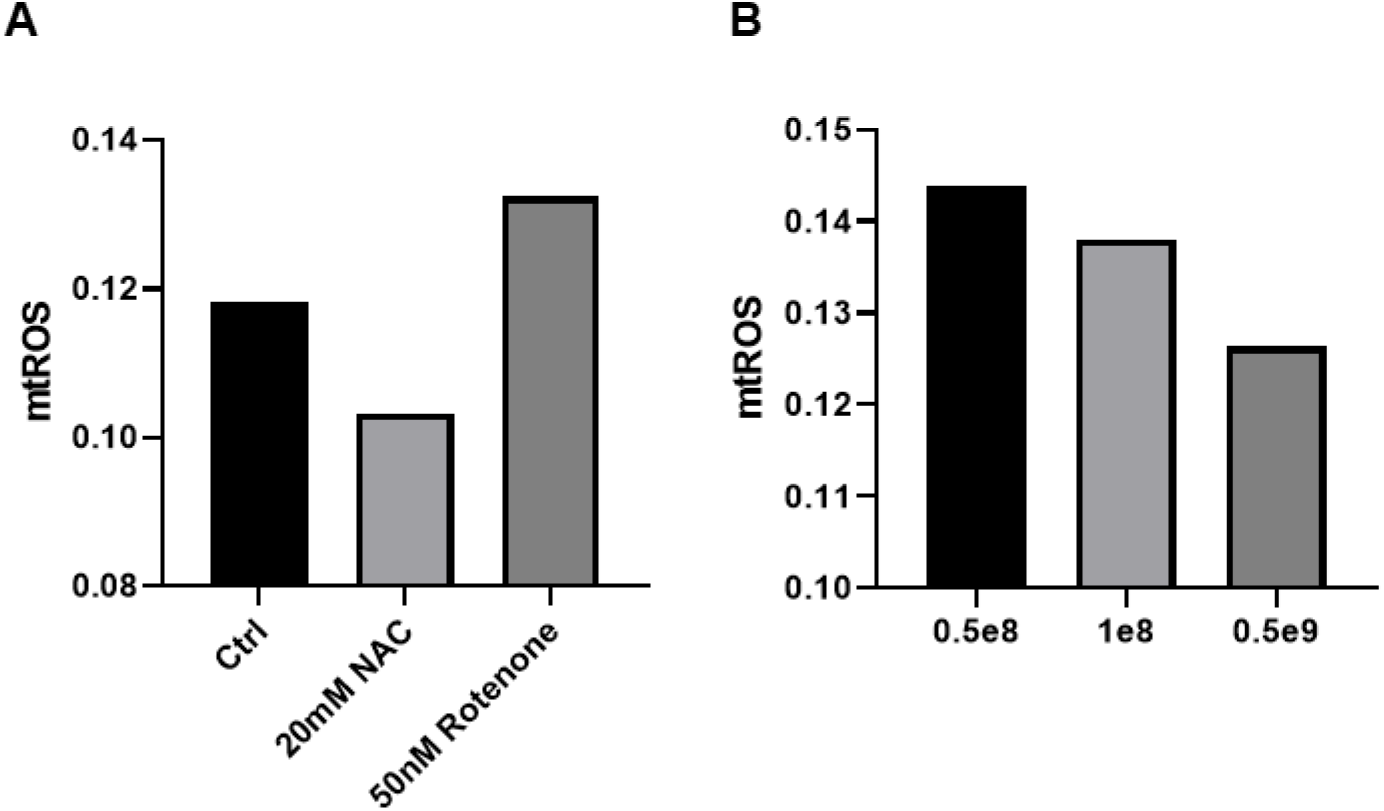
Population average mtROS values after treatment with small molecules or adenoviral SOD2-mitoCAT. (A) Human primary myoblasts treated with DMSO control (Ctrl) or the antioxidant N-acetyl-cysteine (NAC) or the mtROS-inducing drug rotenone. (B) Human primary myoblasts infected with 0.5×10^8^ pfu/ml to 0.5×10^9^ pfu/ml of adenovirus expressing SOD2-mitoCAT.

**Figure S4.**
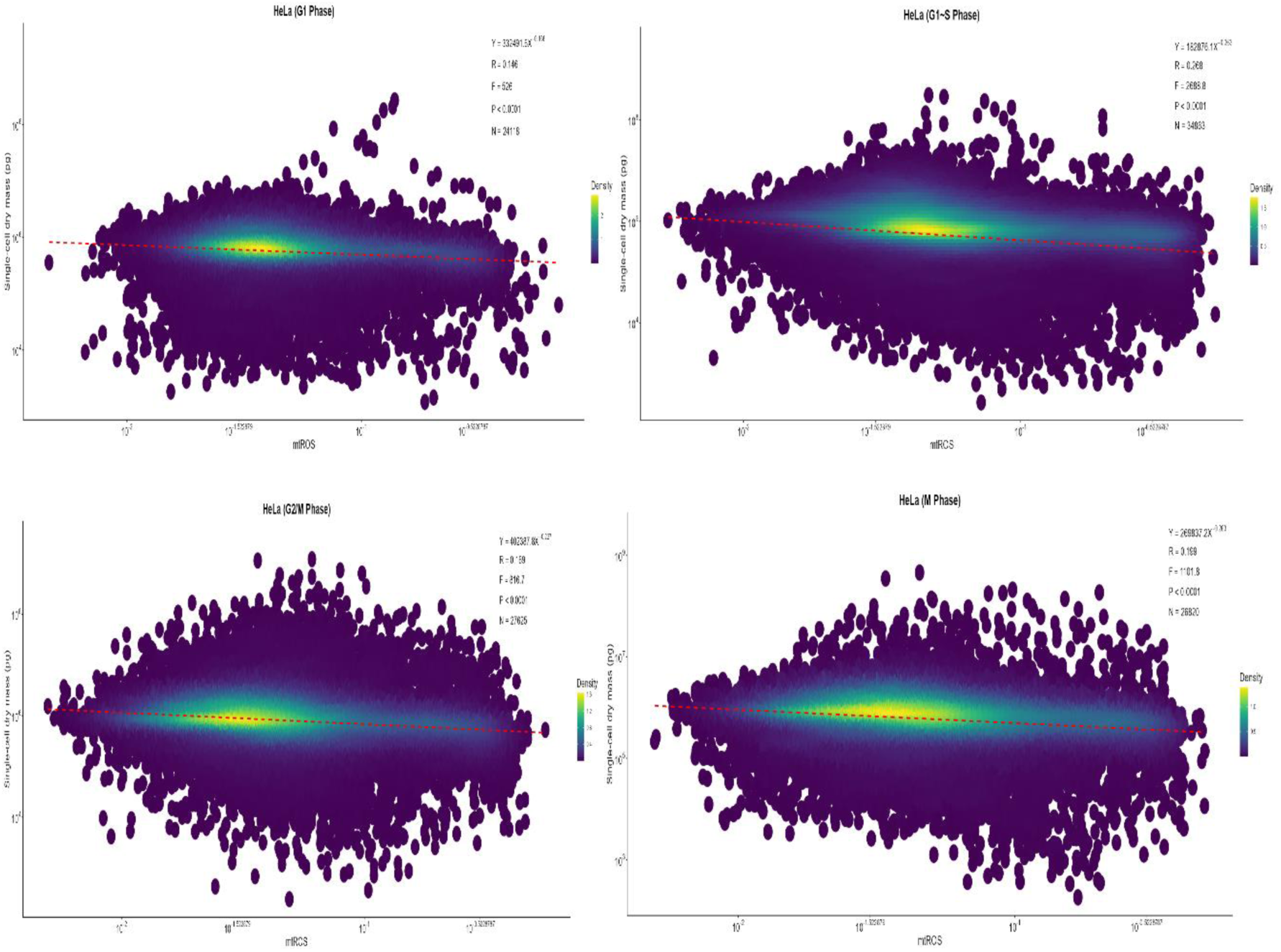
The scMass-mtROS scaling law in synchronized HeLa cells at different stages of the cell cycle.

**Table S1.**
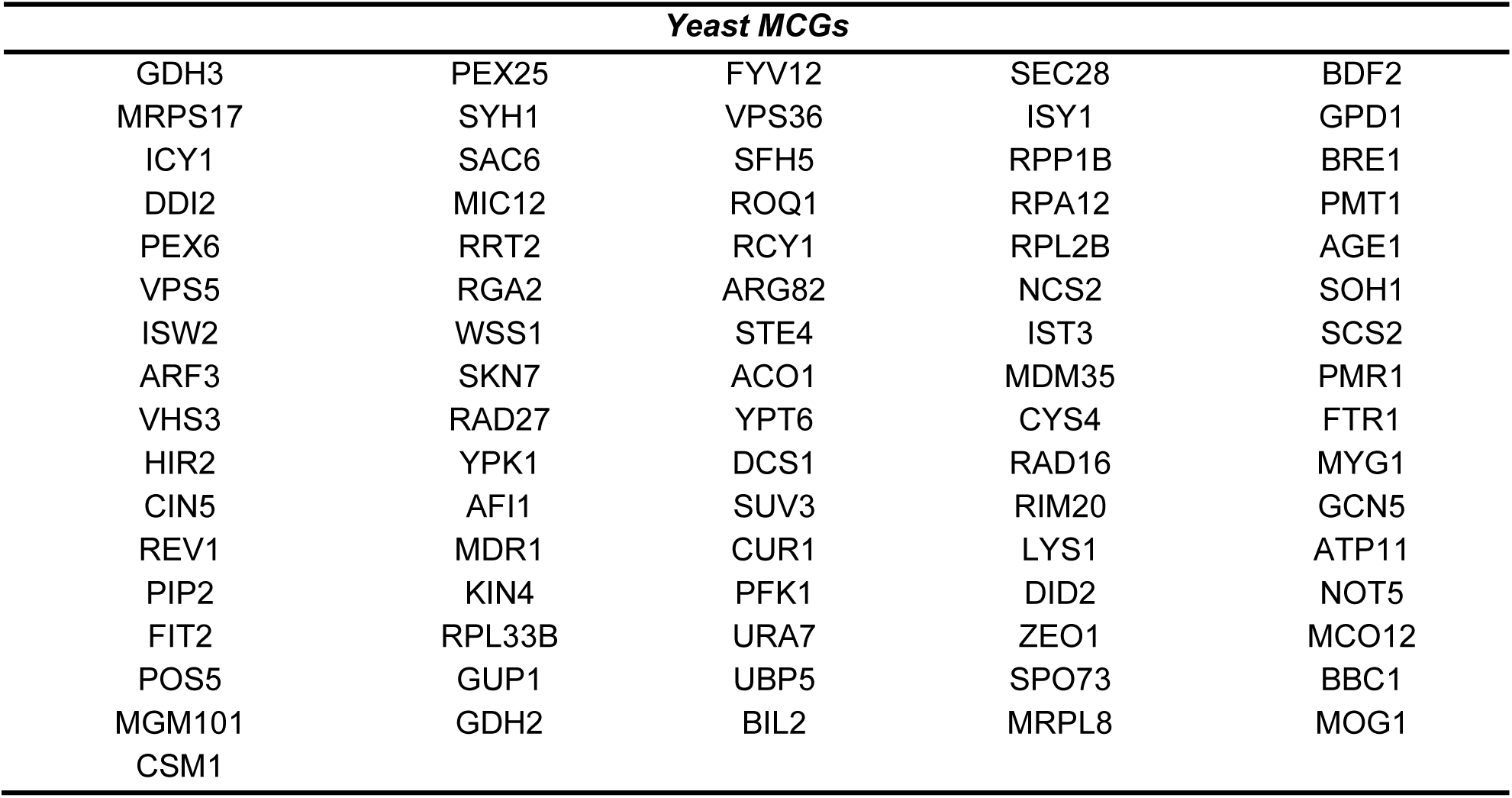
List of MCGs.

## Methods

### Mammalian cell culture

Human cancer cell lines HeLa and HCT116, along with the mouse melanoma cell line B16, were cultured in Dulbecco’s modified Eagle medium (DMEM) supplemented with 10% FBS (BIOIND) and 1% Penicillin-Streptomycin (Gibco) at 37°C and 5% CO_2_. Human adult primary myoblasts were maintained in DMEM with 20% FBS and 1% Penicillin-Streptomycin. Human primary endothelial cells (ECs) were incubated in EBM-2 (Lonza, CC-3156). All cells were passaged at 80%-90% confluency.

### Virus production and cell transduction

The plasmids listed below were utilized to generate viruses for the different transgenic cell lines: pLPCX mito Grx1-roGFP2 (Addgene no, 64977; Gutscher et al., 2008), lentiviral envelope plasmid VSV-G (Addgene no. 8454), PMD2.G (Addgene no, 12259), packaging plasmid dR8.2 (Addgene no. 8455), psPAX2 (Addgene no, 12260), SOD2-TA-mitoCAT (Addgene no, 67635). 293FT cells were seeded at approximately 40% confluency. Mix the aforementioned plasmids with PEI (Lablead) at a 1:3 ratio and add the mixture to the 293FT cells in a dropwise manner. Viral supernatants were harvested within the 48 to 72- hour timeframe and filtered using a 0.45 μm filter (Sartorius). Add the obtained viral supernatants along with polybrene (Lablead) to the corresponding cells. After 3 days, the virally transduced cells were selected with the appropriate antibiotics, typically achieving a transduction efficiency of 80%.

### Yeast cell culture and transformation

The budding yeast used in this study is S. cerevisiae By4742, generously provided by Professor Wei Li. Cells were grown in YPD liquid medium (Solarbio) (OD600 o = 0.5-0.8), then collected and transformed with the pGAP-ATP9-roGFP2-Ura3 plasmid (Miaoling Biotechnology, Wuhan, China) using the lithium acetate method. The transformed cells were carefully plated on SC-URA (OEM) plates and incubated at 30°C for 48 hours to select for positive clones. The budding yeast knockout collection (YKO) library, developed by the Saccharomyces Genome Deletion Project, was acquired from Horizon Discovery (Dharmacon).

### High-throughput imaging

All cells excluding yeast cells were seeded in gelatin-coated 12-well plates at 50% confluency and maintained with culture medium. For yeast cells, transfer 2μL of the cell suspension to a microscope slide, and cover with a coverslip. Images were acquired with a PerkinElmer Operetta CLS High Content Screening System with a ×10 high NA objective. The whole cell images were analyzed for 405, 488 and Digital Phase Contrast (DPC) signals to determine mtROS and cell masses respectively. Images were analyzed by using Harmony 4.6 software. Specifically, cells were extracted and calculated by optimizing for specific parameters, including the cell area and DPC intensity.

### Drug/virus treatment

Human myoblasts were seeded in gelatin-coated 12-well plates at 50% confluency and maintained with culture medium. One day after seeding, culture medium was replaced with culture medium containing either N-acetyl-cysteine (Sigma-Aldrich; 20 mM), Rotenone (MCE; 50 uM) or adenoviral-SOD2-TA-mitoCAT (Vigene) at various concentration gradients.

### Cell dry mass

After digesting the cells used for imaging with 0.25% trypsin and centrifuging, cells were washed twice with PBS and centrifuged at 1000 rpm for 5 minutes. The supernatant was carefully aspirated, leaving only the cell pellet. The wet mass of the cells was measured using a precision microbalance (Sartorius). The cell pellets were then placed in a 65°C drying oven until fully dried overnight, and the dry mass was measured.

### Statistics

All statistical analyses were performed using GraphPad Prism 8.3 and R 4.4.1. P < 0.05 was considered significant. Log-normality was established with the Shapiro-Wilk test. The number of biological (non-technical) replicates for each experiment is indicated in the legends.

### Declarations

The authors declare there are no conflicts of interests/competing interests. No biological materials or ethics approvals were involved in this research. All data will be made publicly available before publication.

### Funding Declaration/Acknowledgements

This work is supported by the Howard Hughes Medical Institute International Scholar grant, and the National Key R&D Program of China (Grant No. 2019YFA0801700).

